# Knockdown resistance (kdr) Associated organochlorine Resistance in mosquito-borne diseases (*Culex quinquefasciatus*): Systematic study of reviews and meta-analysis

**DOI:** 10.1101/2024.02.13.580052

**Authors:** Ebrahim Abbasi, Salman Daliri

## Abstract

**Introduction:** *Culex quinquefasciatus* is one of the most important carriers of human pathogens. The use of insecticides is one of the most important methods of combating this vector. But the genetic resistance created in *Culex quinquefasciatus* has led to disruption in the fight against this pest. As a result, it is necessary to know the level of resistance in order to fight this vector. Based on this, the present study was conducted with the aim of investigating the prevalence of kdr resistance in *Culex quinquefasciatus* against organochlorine insecticides in the world.

**Methods:** This study was conducted by systematic review and meta-analysis on the prevalence of kdr resistance and mortality rate in *Culex quinquefasciatus* against organochlorine insecticides in the world. Based on this, during the search in the scientific databases Web of Science, PubMed, Scopus, biooan.org, Embase, ProQuest and Google scholar without time limit until the end of November 2023, all related articles were extracted and analyzed. Statistical analysis of the data was done using fixed and random effects model in meta-analysis, *I^2^* index, Cochran’s test and meta-regression by STATA version 17 software.

**Results:** Seventy articles were included in the meta-analysis process. Based on the findings, the prevalence of Kdr in *Culex quinquefasciatus* against organochlorine insecticide was estimated at 63.1%. Also, the mortality rate against the insecticide deltamethrin was 46%, DDT 18.5%, permethrin 42.6%, malathion 54.4% and lambdacyhalothrin 53%.

**Conclusion:** More than half of Cx. quinquefasciatus had Kdr. This vector was relatively resistant to DDT and permethrin insecticides and sensitive to malathion, deltamethrin and lambdacyhalothrin. As a result, it is necessary to use effective insecticides to fight this vector in order to prevent the increase of resistance to other insecticides.

## Introduction

Mosquitoes carry various types of diseases, including malaria, dengue fever, West Nile fever, yellow fever, filariasis, Japanese encephalitis, and Zika, especially in tropical and subtropical regions of the world. The genus Culex is one of the types of mosquitoes and has 550 species. *Culex quinquefasciatus* species, *Cx. fuscocephala, Cx. pseudovishnui, Cx. gelides, Cx. tritaenorhynchus* and *Cx. vishnui* are carriers of diseases including Japanese encephalitis, Bancroftian filariasis, West Nile virus, and St. Louis encephalitis virus (1). *Culex quinquefasciatus* is the major vector of Bancroftian filariasis in Asia, South Africa and India (1). Due to the endemicity of Bancroftian filariasis in India, about six hundred million people are exposed to this disease (2).

For many decades, the use of synthetic insecticides to control vectors has been the most important measure in the field of managing vector-borne diseases. As a result, people used insecticides in the form of indoor residual sprays (IRS), insecticide-impregnated nets (ITNs), and LLINs to control and fight these vectors and prevent the spread of diseases. But the widespread and indiscriminate use of chemical insecticides over time led to resistance in the vector population (3, 4).

Resistance developed in insects against insecticides is carried out through four mechanisms, which include 1) modifying or changing behavior 2) reducing the penetration of insecticides into the body, by increasing the thickness of the cuticle or changing the composition of the cuticle 3) metabolic detoxification of the insecticide which is done by strengthening or changing key enzymes. 4-Mutation in the target location and reducing their sensitivity (5, 6). Mutations in the target site and metabolic detoxification by enzymes are among the most important and widespread mechanisms of resistance to chemical insecticides (7). Insensitivity to the target site is known as knockdown resistance (kdr), it is caused by a mutation in the voltage-sensitive sodium channel gene (Vssc) and is one of the most common types of resistance in insects (6).

Organophosphates, organochlorines, pyrethroids, and carbamates are the four main groups of insecticides to fight against vectors (8). Organochlorines are widely used to control vectors including *Culex quinquefasciatus*. These insecticides affect the central and peripheral nervous system of insects and through the effect on the sodium channel and increasing its sensitivity to depolarization along with inhibiting the inactivation processes, lead to paralysis and death of the vectors (9, 10). This type of resistance leads to the neutralization of DDT and Deltamethrin to fight against vectors and reduce their nervous sensitivity to these insecticides (11).

*Cx. quinquefasciatus* resistance against insecticides has been reported from different regions of the world and is increasing (12, 13). Different studies have shown the presence of kdr resistance and reduced effect of organochlorine and organophosphorus insecticide in *Cx. quinquefasciatus* have reported, but the prevalence of these resistances has been different in different regions of the world (14, 15). Based on this, in order to fight this vector, it is necessary to know the resistance and choose the right insecticide. Considering the spread of this vector and the diseases caused by it in the world, this study was carried out with the aim of the prevalence of Kdr and the mortality rate against organochlorine insecticides in the world by systematic review and meta-analysis.

## Materials and Methods

This systematic review and meta-analysis study was conducted on the prevalence of kdr in *Cx. quinquefasciatus* and resistance to organochlorine insecticides based on the guidelines of the Preferred Reporting Items for Systematic Reviews and Meta-Analyses (PRISMA) (16). This research has been registered in the International Prospective Register of Systematic Review (PROSPERO) with the code CRD42021231605.

### Search Strategy

By searching in scientific databases Web of Science, PubMed, Embase, ProQuest, biooan.org, Scopus, and Google Scholar and using the keywords knockdown resistance, Resistance, KDR, Organochlorine Insecticide, Insecticide, Chlorinated Insecticide, Chlorophenyl, DDT, malathion, dichloroethane, para chlorophenyl, dichlorodiphenyldichloroethane, dieldrin, deltamethrin, permethrin, *Culex*, *Culex quinquefasciatus*, all related articles were extracted without time limit until the end of 2023. Searching for keywords was done individually and in combination using OR, AND, and NOT operators in the title, abstract, and full text of the articles.

### Inclusion And Exclusion Criteria

The inclusion criteria included 1-English language articles that were conducted on *Cx. quinquefasciatus*, 2-Prevalence of resistance or mortality in exposure to organochlorine insecticides was reported or estimated in them, 3-KDR resistance was investigated in them., 4- and they had good quality. Exclusion criteria included: conducting a study on other insects, not investigating kdr resistance or mortality, not having the desired quality and conducting the study using a qualitative method, reporting cases or cases, a review or a narrative, and a letter to the editor.

### Quality Assessment

Using the STROBE (Strengthening the Reporting of Observational Studies in Epidemiology) checklist, the quality of the articles was measured. This checklist in the field of observing the principles of research writing and implementation has the areas of title, project implementation, research findings, limitations, and conclusions. that each part of this checklist has subgroups and points are given based on their importance. The maximum score that can be obtained is 33 and the obtained score is more than 20 is acceptable (17).

### Data Extraction

At first, taking into account the inclusion and exclusion criteria, the title and abstract of all articles were reviewed by two researchers independently. If the articles were related, the full text was checked and if they were unrelated, they were excluded from the study. In cases where there was a difference of opinion between two researchers, the article was refereed by a third person. Data extraction was done using a pre-prepared checklist that included the first author’s name, year of study, study location, sample size, insecticide type, KDR resistance prevalence, and mortality rate.

### Selection of Studies

the number of 16,852 articles extracted from scientific databases were entered into Endnote software. At first, duplicate articles were reviewed, and 7654 articles were removed due to duplicates. In the next step, after reviewing the title, abstract, and the full text of the articles, 9185 articles were removed due to being unrelated or not investigating kdr resistance, resistance to organochlorine insecticide or mortality rate, and finally 13 articles met the inclusion criteria and entered the meta-analysis process (Figure 1).

**Figure 1.**
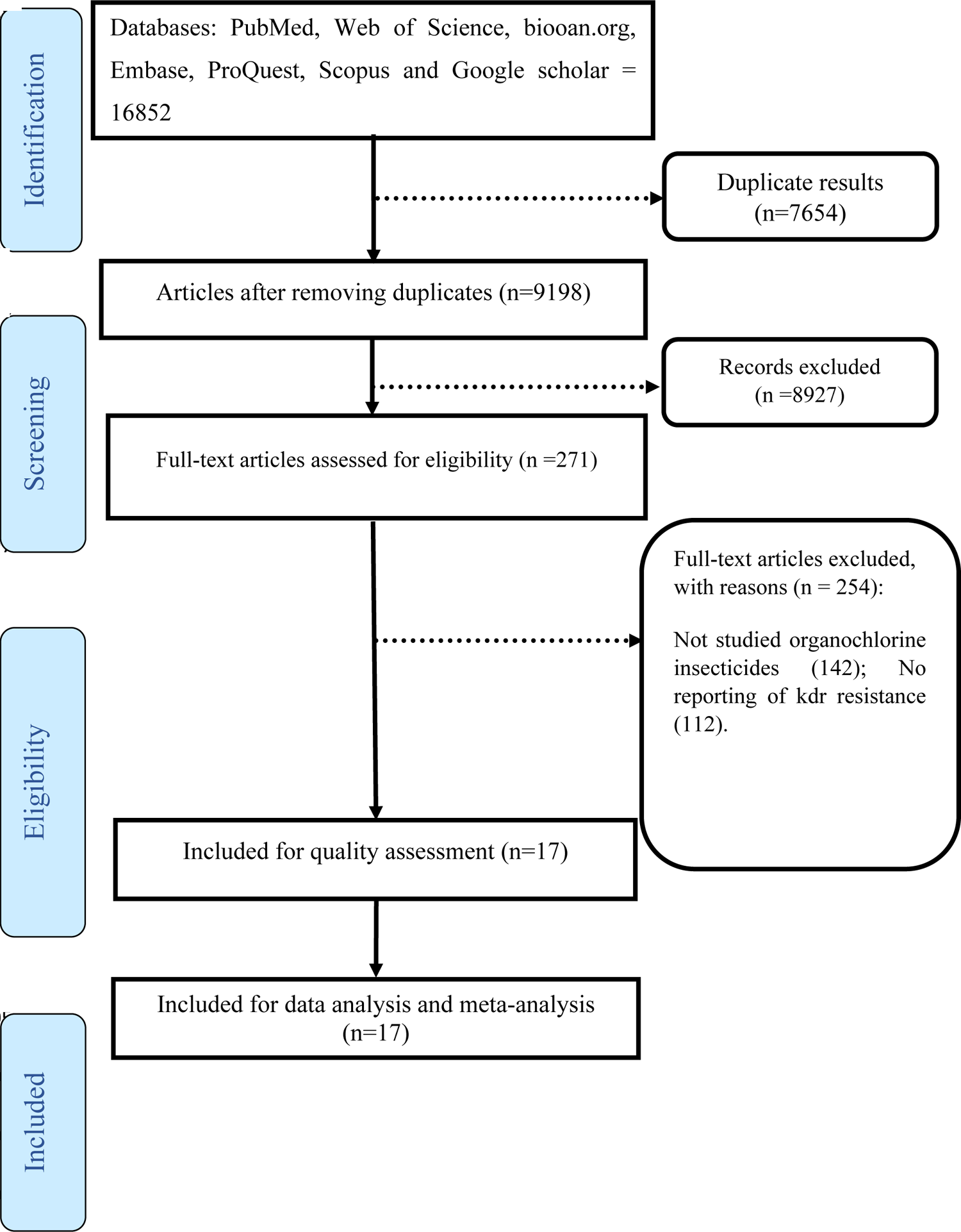
The PRISMA flow diagram

### Statistical Analysis

Data were analyzed using fixed and random effects models in the meta-analysis, Cochran test, and I2 index. Egger’s test and funnel plot were used to check the publication bias and meta-regression was used to check the relationship between KDR prevalence and sample size. Data were analyzed using STATA version 17 software.

## Results

seventy articles with a sample size of 3733 that were conducted between 2003 and 2020 were included in the study. 13 articles investigated the prevalence of Kdr resistance. Also, the mortality rate against deltamethrin was investigated in 13 articles, against DDT in 11 articles, against permethrin in 11 articles, against malathion in 3 articles, and against lambda-cyhalothrin in 3 articles. The characteristics of the reviewed articles are presented in Table 1. investigate the prevalence of Kdr resistance against organochlorine insecticides showed that 63.1% of *Cx. quinquefasciatus* had Kdr resistance (Figure 2). Also, based on the type of resistance, the prevalence of homozygote resistance was estimated at 28.6%, and the prevalence of heterozygote resistance at 37.4% (Figures 3 and 4).

**Figure 2.**
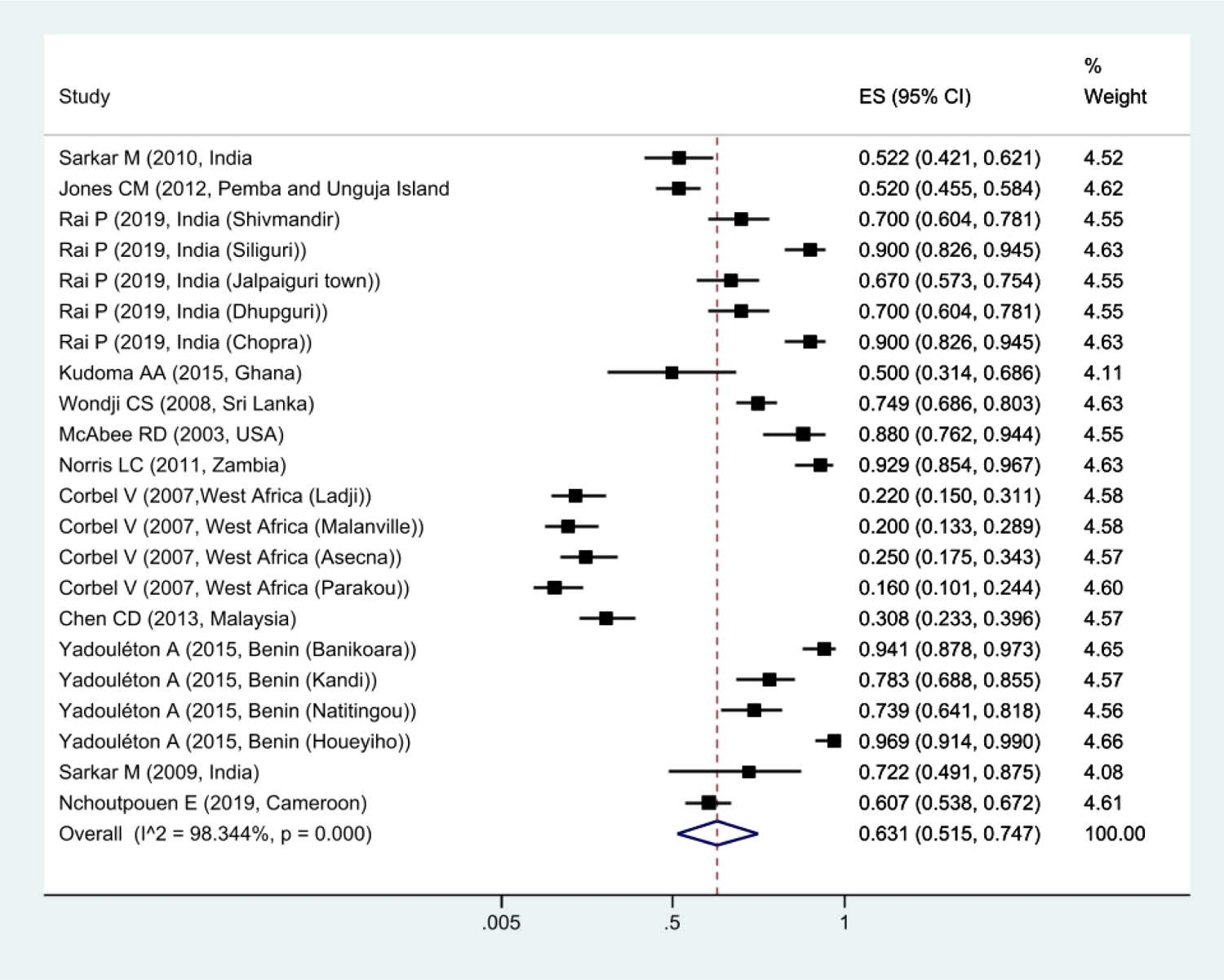
Forest plots of the prevalence Kdr and 95% confidence interval based on the random effect model in meta-analysis.

**Figure 3.**
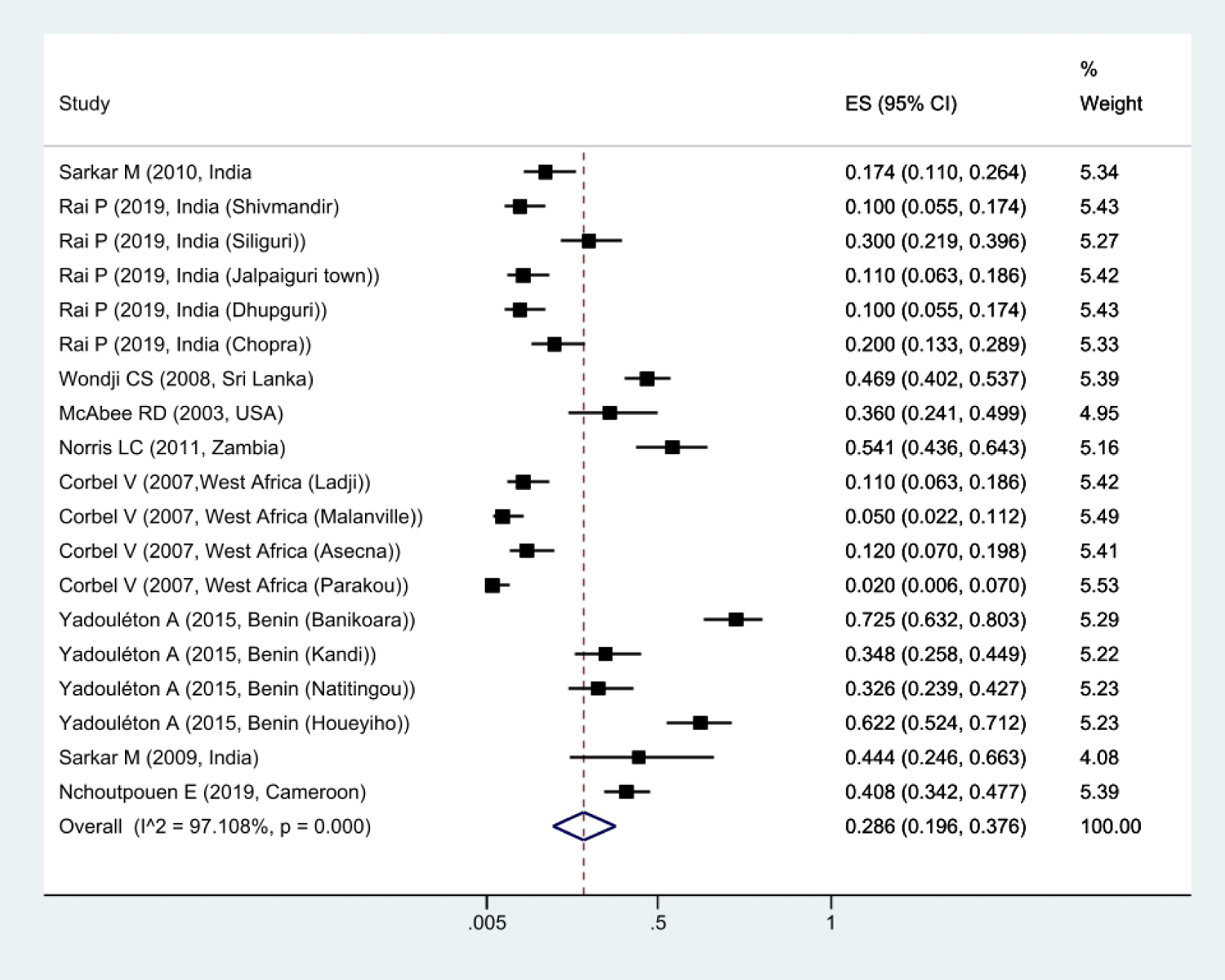
Forest plots of the prevalence homozygotes resistance and 95% confidence interval based on the random effect model in meta-analysis.

**Figure 4.**
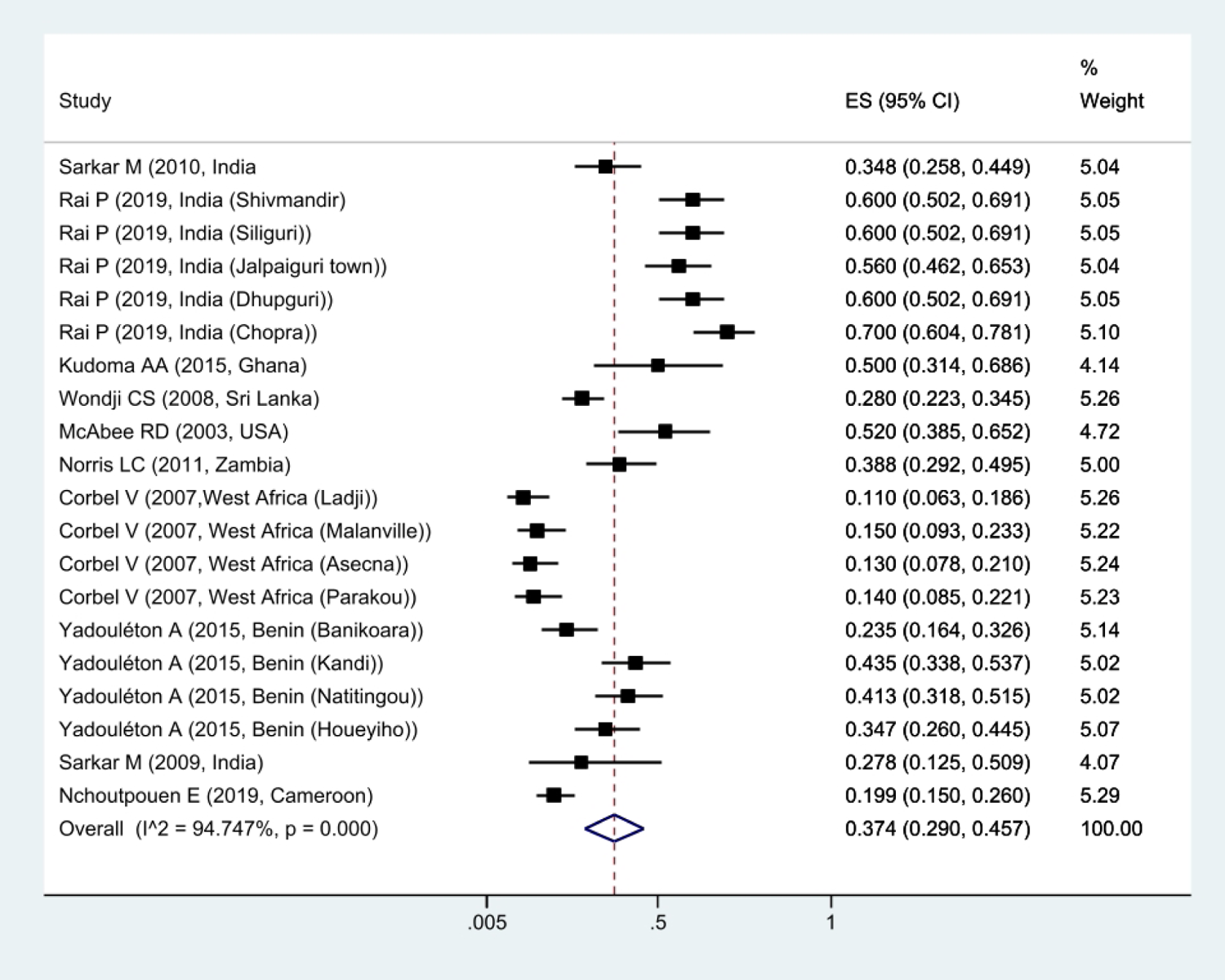
Forest plots of the prevalence heterozygotes resistance and 95% confidence interval based on the random effect model in meta-analysis.

**Table 1.**
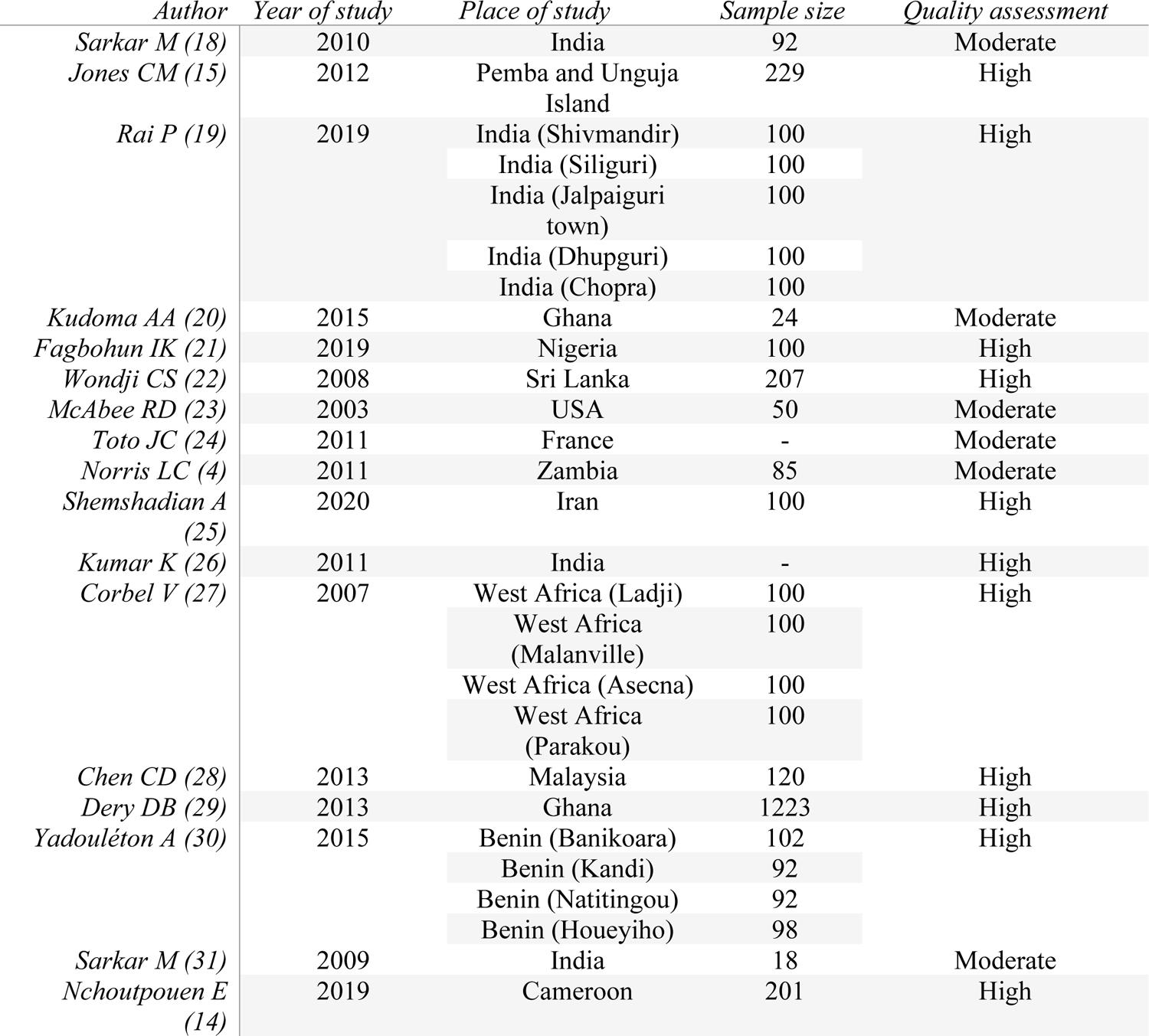
Characteristics of the articles included in the meta-analysis process.

The meta-analysis of the mortality rate based on the type of insecticide used to fight *Cx. quinquefasciatus* showed that the mortality rate was 46% against deltamethrin, 18.5% against DDT, 42.6% against permethrin, and 54.4% against malathion and 53% against lambda-cyhalothrin, it was estimated. This shows that *Cx. quinquefasciatus* was most sensitive to Malathion insecticide and least sensitive to DDT (Figure 5-9).

**Figure 5.**
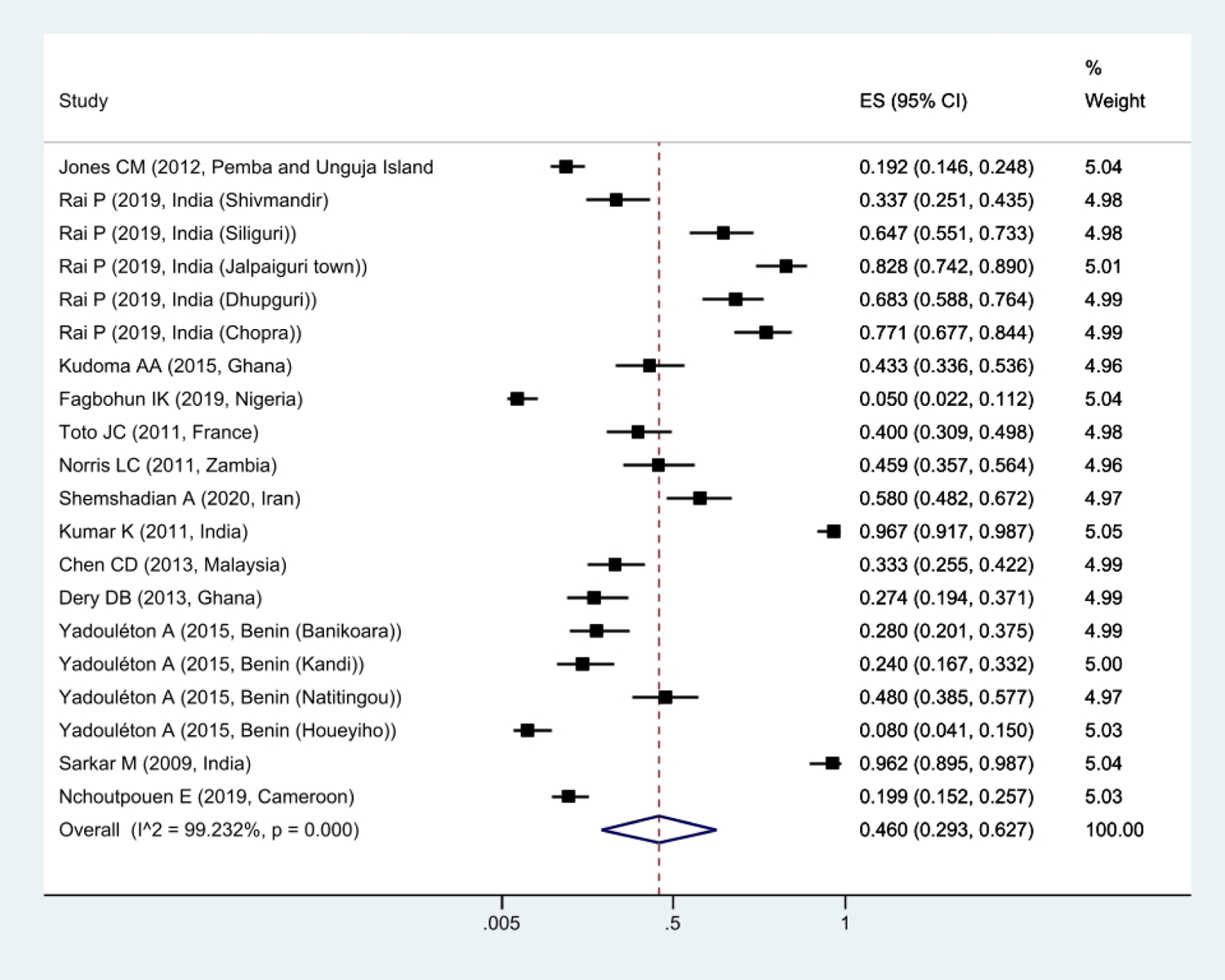
Forest plots of the mortality rate *Cx. quinquefasciatus* exposed to Deltamethrin and 95% confidence interval based on the random effect model in meta-analysis.

**Figure 6.**
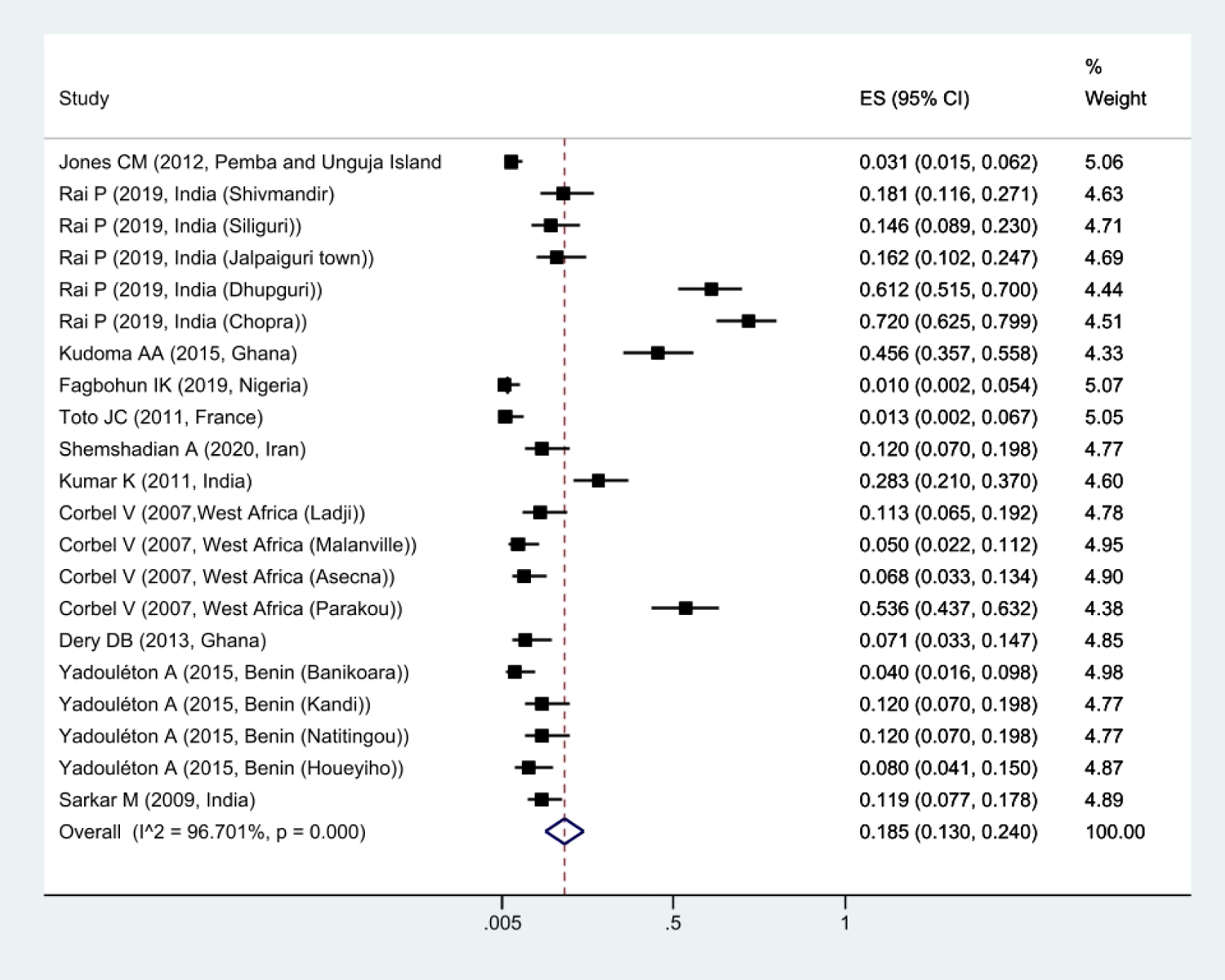
Forest plots of mortality rate *Cx. quinquefasciatus* exposed to DDT and 95% confidence interval based on the random effect model in meta-analysis.

**Figure 7.**
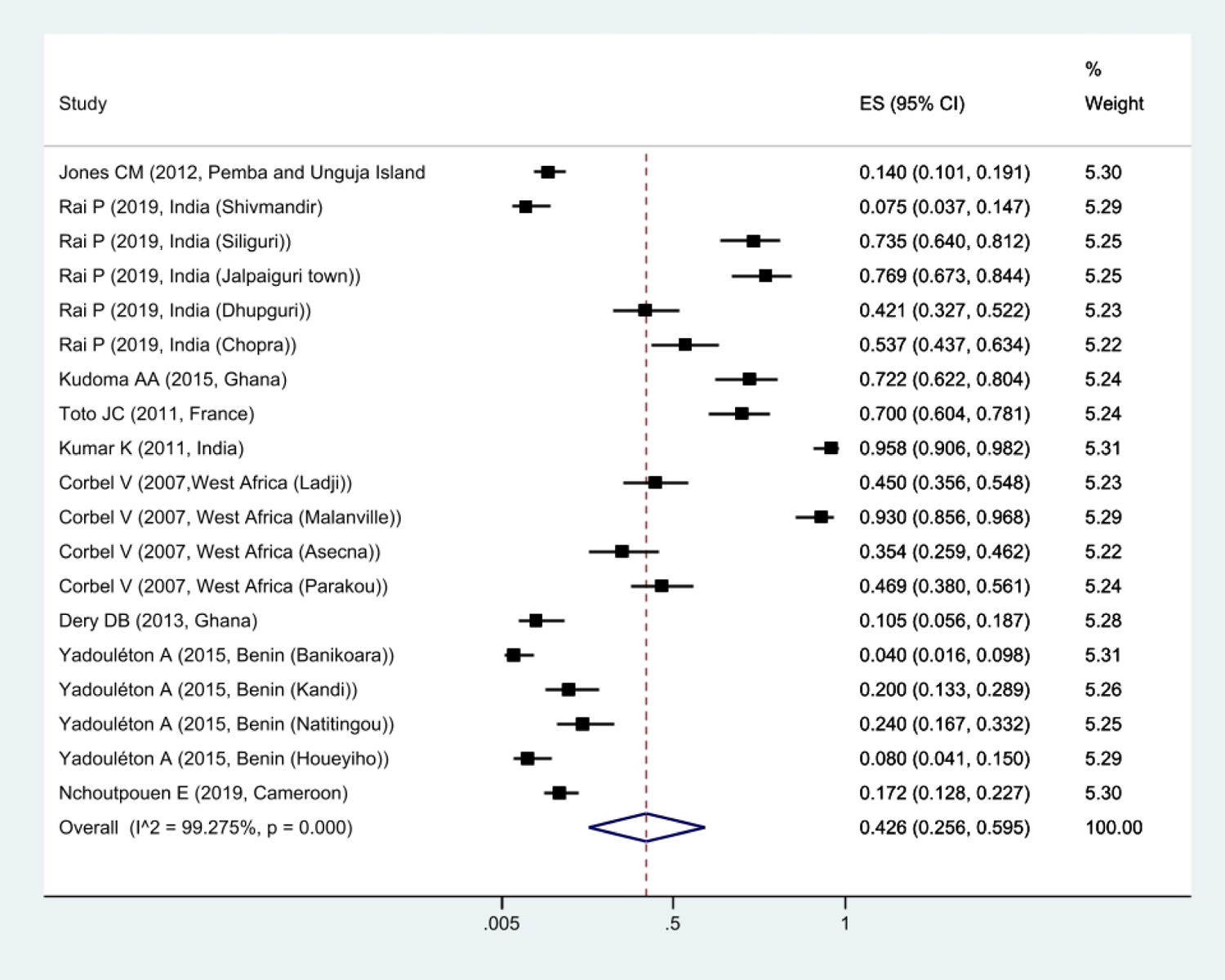
Forest plots of mortality rate *Cx. quinquefasciatus* exposed to Permethrin and 95% confidence interval based on the random effect model in meta-analysis.

**Figure 8.**
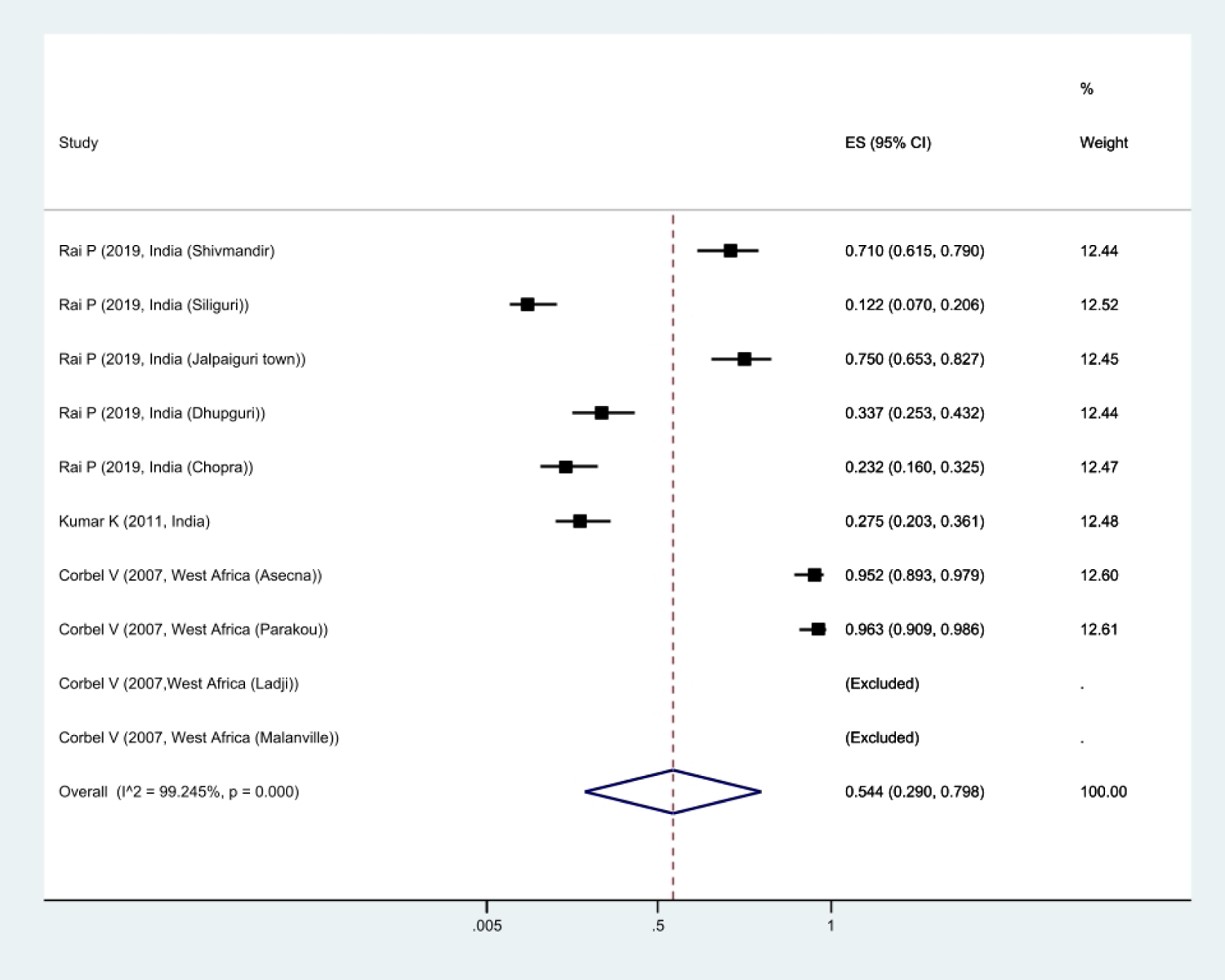
Forest plots of mortality rate *Cx. quinquefasciatus* exposed to Malathion and 95% confidence interval based on the random effect model in meta-analysis.

**Figure 9.**
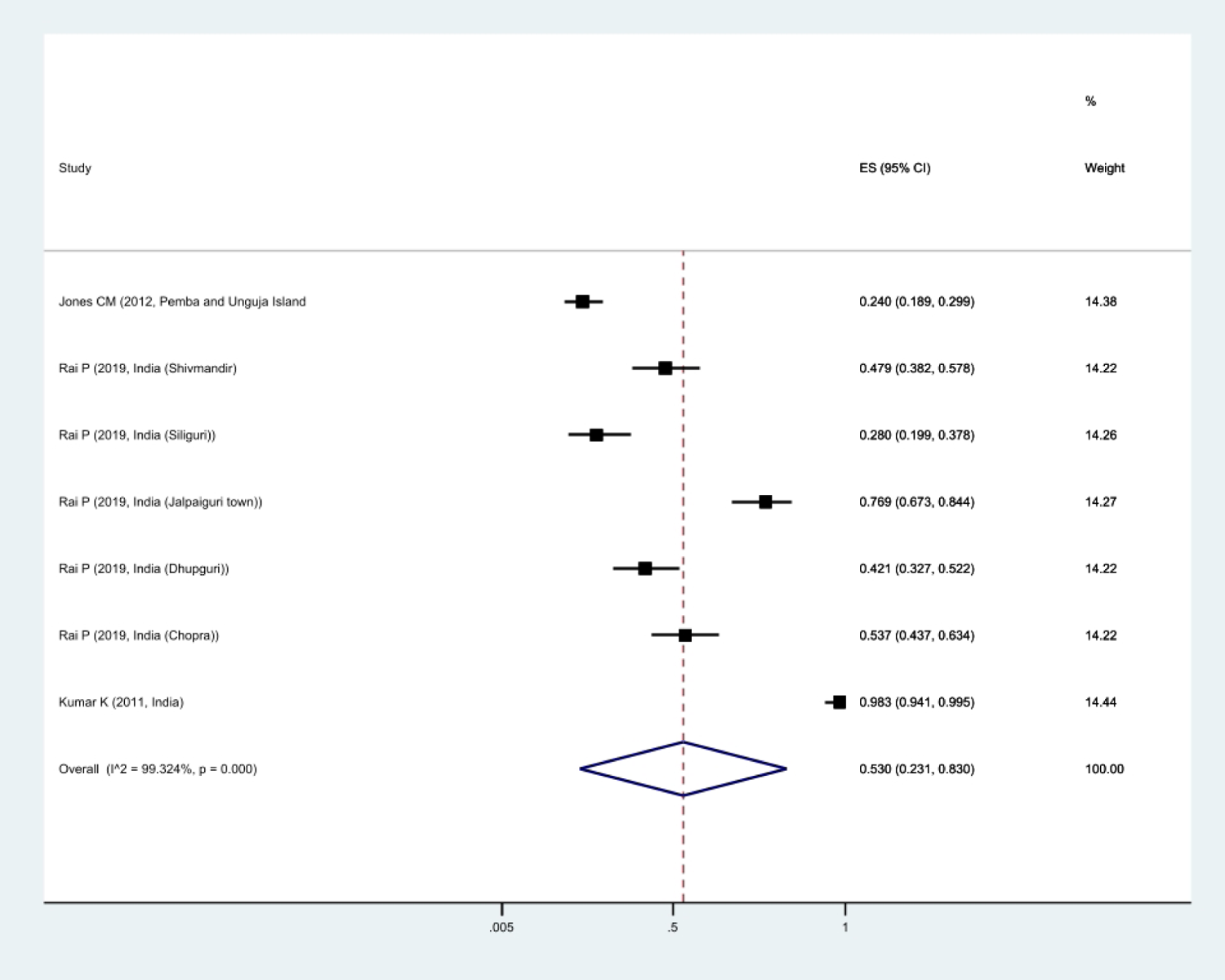
Forest plots of mortality rate *Cx. quinquefasciatus* exposed to lambdacyhalothrin and 95% confidence interval based on the random effect model in meta-analysis.

Publication bias was done using funnel plot, Egger’s test, and investigating the relationship between sample size and mortality rate using meta-regression. Due to the placement of studies with a high sample size below the graph, it can be mentioned that the publication bias did not occur (Figure 10). This result was confirmed due to the non-significance of Egger’s test (P=0.145). According to the slope of the meta-regression graph, with the increase in the sample size, the mortality rate decreased (Figure 11).

**Figure 10.**
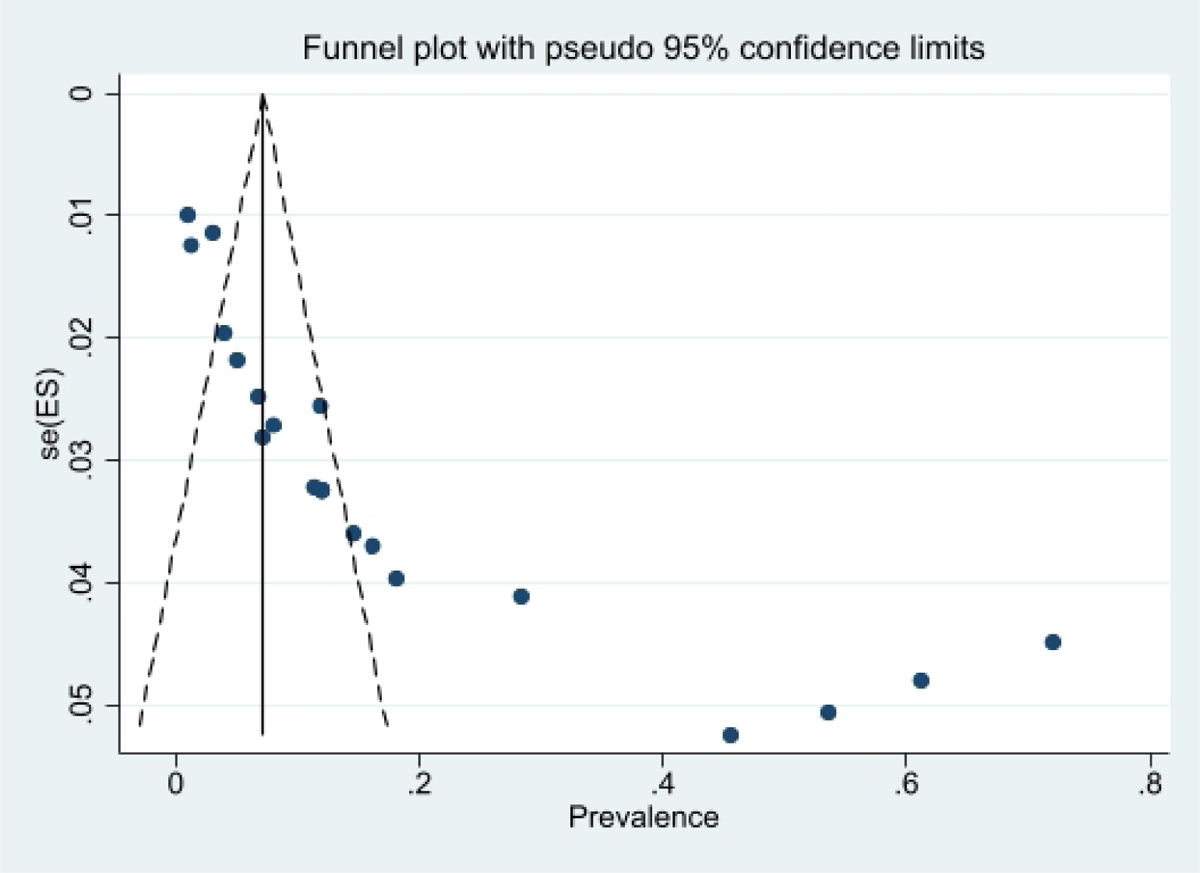
Funnel plot of the mortality rate *Cx. quinquefasciatus* in the selected studies

**Figure 11.**
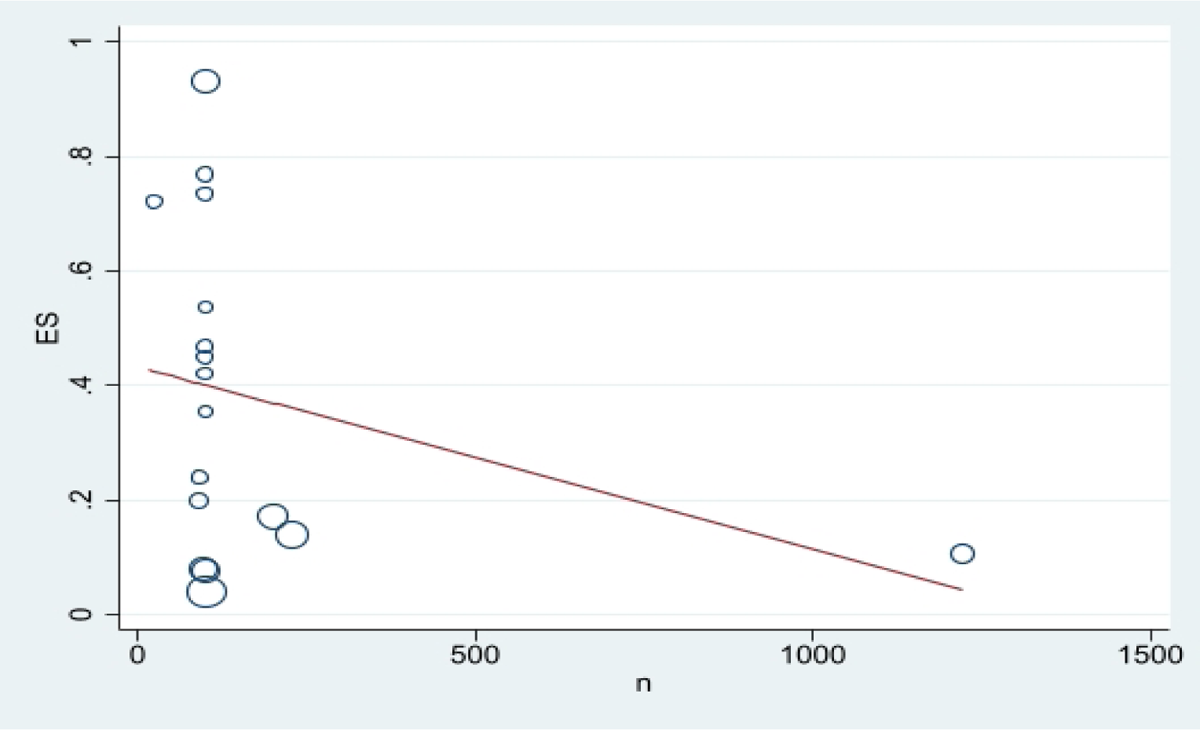
Meta regression plot of the mortality rate *Cx. quinquefasciatus* of exposed to Permethrin based the study year.

## Discussion

In the present study, the prevalence of Kdr and its types in *Cx. quinquefasciatus*, as well as the mortality rate against permethrin, DDT, malathion, deltamethrin, and lambda-cyhalothrin insecticides, were investigated. Based on the results of the meta-analysis, more than half of *Cx. quinquefasciatus* had Kdr against organochlorine insecticides. In the field of Kdr resistance, the prevalence of resistance heterozygotes was higher. Kdr mutations are the most important resistance mechanism of insects against pyrethroids and DDT(9). Various studies have reported the role of kdr mutation in the resistance of *Cx. quinquefasciatus* against DDT and permethrin insecticides (32–34). The most important mechanism of Kdr resistance in this transporter is the substitution of L to F at position 1014. This substitution is involved in high resistance to DDT and pyrethroids (35, 36). In Xu et al.’s (2005) study, heterozygous and homozygous mutations for the kdr allele in Cx. quinquefasciatus, it was observed that the frequency of heterozygous was much higher than homozygous (34). In Xu et al.’s (2006) study, the relationship between the kdr allele at the genomic DNA level and the susceptibility and resistance of *Cx. quinquefasciatus* was not related to insecticides. However, a strong correlation between kdr allelic expression through RNA allelic diversity and the level of resistance and sensitivity to insecticides was observed (37). In general, studies conducted in different regions of the world have presented various reports on the prevalence of Kdr and its role in the field of resistance to insecticides. According to the present study, more than half of *Cx. quinquefasciatus* have Kdr resistance against organochlorine insecticides, and this mutation has led to high resistance against these insecticides. Considering that these mutations can be exchanged between insects, it can lead to increased resistance to other insecticides. As a result, it is recommended before using insecticides to fight *Cx. quinquefasciatus* Kdr should be checked them and the appropriate insecticide should be used.

In the present study, high resistance in *Cx. quinquefasciatus* was observed against DDT and then permethrin. In the conducted studies, it was observed that mosquitoes resistant to DDT were also resistant to pyrethroid insecticides over time. The reason can be the indiscriminate use of pyrethroids and selection pressure, which ultimately leads to the ineffectiveness of vector control activities and pyrethroid insecticides (38, 39). Studies have shown that mosquitoes that were previously resistant to DDT can also become resistant to pyrethroids due to cross-resistance. Resistance to DDT and pyrethroids has caused a major concern in the field of vector control, especially their use for IRS and LLIN, which are important tools for vector control (40, 41). In Low et al.’s (2013) study, adult mosquitoes of *Cx. quinquefasciatus* were highly resistant to permethrin and DDT. However, the larvae of *Cx. quinquefasciatus* were relatively more resistant to malathion and sensitive to permethrin (42). The difference in the level of sensitivity to insecticides between larvae and adult mosquitoes is due to the difference in the expression of the resistance gene in the larval and adult stages. Based on studies conducted on resistance to insecticides in the larval stage of *Cx. quinquefasciatus* is more common than adult mosquitoes. The reason can be excessive use of insecticides and selection pressure (43–45). Studies conducted on populations of *Cx. quinquefasciatus* have shown that resistance to insecticides in this vector is correlated with each other, which indicates cross-resistance, although their mechanism is unknown (46). In general, cross-resistance between pyrethroids, organochlorines, organophosphorus, and carbamates has been observed in vectors and it has been mentioned that high levels of functional oxidases can cause cross-resistance in insecticides (47, 48). In Nazni et al.’s (2005) study, it was noted that DDT was the least effective insecticide among all insecticides tested for *Cx. quinquefasciatus* (49).

In general, the bioassay of *Cx. quinquefasciatus* is necessary to evaluate resistance to insecticides and to know the ratio of resistance to combat this vector. In the present study, *Cx. quinquefasciatus* had high resistance to DDT and permethrin and relatively high sensitivity to malathion, deltamethrin, and lambdacyhalothrin. that these insecticides can be used as effective insecticides to fight against it. Of course, it should be mentioned that the prevalence of resistance is different in different regions and it is recommended to check the resistance before choosing insecticides for the fight.

## Conclusion

According to the findings, more than half of *Cx. quinquefasciatus* had Kdr. *Cx. quinquefasciatus* had relatively high resistance to DDT and permethrin insecticides and was sensitive to malathion, deltamethrin, and lambda-cyhalothrin. Based on this, we can mention these insecticides to fight *Cx. quinquefasciatus* are effective. However, it is recommended to first evaluate the resistance of these insecticides to determine their effectiveness.

## Declaration

### Ethics approval and consent to participate

Not applicable.

### Availability of data and materials

All data obtained from this study are included in the text of article.

### Authors’ contributions

EA determined the title, wrote and registered the protocol, and submitted the article. EA and SD extracted the files from the databases. EA and SD, screening, and selection of final reports. EA, data extraction. SD wrote the article. All authors read and approved the final manuscript.

